# Cell type-dependent control of p53 transcription and enhancer activity by p63

**DOI:** 10.1101/268649

**Authors:** Gizem Karsli Uzunbas, Faraz Ahmed, Morgan A. Sammons

**Affiliations:** Department of Biological Sciences, State University of New York at Albany, Albany, NY 12222

## Abstract

Transcriptional activation by p53 provides powerful, organism-wide tumor suppression. In this work, we demonstrate that the p53-induced transcriptome varies based on cell type, reflects cell type-specific activities, and is considerably more broad than previously anticipated. This behavior is strongly influenced by p53 engagement with differentially active cell type-specific enhancers and promoters. In epithelial cell types, p53 activity is dependent on the p53 family member p63, which displays widespread enhancer binding. Notably, we demonstrate that p63 is required for epithelial enhancer identity including enhancers used by p53 during stress-dependent signaling. Loss of p63, but not p53, leads to site-specific depletion of enhancer-associated chromatin modifications, suggesting that p63 functions as an enhancer maintenance factor in epithelial cells. Additionally, a subset of epithelial-specific enhancers is dependent on the activity of p63 providing a direct link between lineage determination and enhancer structure. These data suggest a broad, cell-intrinsic mechanism for regulating the p53-dependent cellular response to stress through differential regulation of *cis*-regulatory elements.

## INTRODUCTION

The p53 family of transcription factors regulate highly diverse cellular functions, including tumor suppression and control of cell specification and identity (1). p53 is a master tumor suppressor that protects organismal fidelity after exposure to cellular stress like DNA damage. This activity is dependent on p53’s ability to activate transcription of a canonical network of genes involved in DNA damage repair, cell cycle arrest, apoptosis, and senescence (2). Surprisingly, many of these transcriptional pathways are individually dispensable for tumor suppression, suggesting p53 regulates a redundant and not-yet fully characterized transcriptional network (3, 4).

Genetic inactivation of the p53 tumor suppressor pathway is highly recurrent across cancer types. p53 mutations, while frequent, vary depending on the tumor type with additional genetic and epigenetic mechanisms proposed to inactivate the p53 pathway in the presence of wild-type p53 (5–7). p63 and p73, p53 family members, function similarly to p53 in stress response, although their precise roles in tumor suppression are unresolved (8–10). p63 and p73 are primarily lineage restricted to epithelial cell types where each serves critical and non-overlapping roles in cell identity and self-renewal (1, 11). Mutations in TA and ΔNp63 isoforms of p63 are causative for a number of epithelial-associated human developmental disorders independent of p53 activity, and mutations in p63 target genes underlie similar phenotypes (12, 13). A significant and still outstanding question involves dissection of specific roles and functional interplay between p53 family members during development, in the regulation of cellular homeostasis, and in the etiology of disease.

The similarity between DNA binding motifs suggested that competition for binding sites serves a central role in the regulation of p53 family member activity (14–16). Indeed, previous studies implicate the ΔNp63 isoforms, lacking transactivation domains, as direct repressors of the p53-induced transcriptome through a binding site competitive mechanism (17–19). Amplification of p63 is thought to drive a number of epithelial cancers, particularly squamous cell carcinomas, although both p53-dependent and independent mechanisms have been proposed for this driver activity (20). As the majority of cancers are derived from epithelial tissues, the mechanisms of p53 family dependent tumor suppression in those tissues is of special interest (7).

Fate choice after p53 activation, be it apoptosis, temporary/permanent cell cycle arrest, or continued proliferation, is variable across cell types suggesting that p53- dependent transcription is also cell type-dependent (2). A bevy of sensitive methodological approaches have been used to identify p53 binding sites and gene targets across transformed and primary cell lines in an effort to explain these terminal cell fate choices and p53-dependent tumor suppression. A recent set of meta-analyses have suggested that p53 binding to the genome is invariant (21), proposing that p53 acts independently to drive gene expression of a core tumor suppressor network across all cell types due to the low enrichment of other transcription factor motifs at p53-bound enhancers and the reported pioneer factor activity of p53 (3, 21, 22). Conversely, similar experimental approaches demonstrate cell type-specific p53 binding and gene targets although the mechanisms driving these differential activities are unknown (BioRxiv: https://doi.org/10.1101/177667). Additional experimental evidence and models are required to unravel these mutually exclusive p53 regulatory mechanisms.

We previously proposed that the local chromatin environment, including variable chromatin accessibility and enhancer activity, contributes to novel p53 activities across cell types (23). To address this question, we performed genomewide transcriptome, epigenome, and p53 cistrome profiling in primary foreskin fibroblasts (SkFib) and breast epithelial (MCF10A) cell lines, two cell types with varying enhancer activity at p53 binding sites (23). Our results directly implicate differential *cis*-regulatory element activity as a mediator of the p53 network, with both promoter and enhancer activity contributing to p53-dependent gene expression differences. Further, we have identified the p53 family member p63 as one factor that drives the epithelial-specific p53 transcriptome through an enhancer maintenance activity. We further propose that p63 serves as licensing factor for a set of epithelial-specific enhancers. Thus, these data support a mechanism whereby co-operating transcription factors control cell type-dependent *cis-*regulatory networks that regulate p53 activity.

## MATERIALS AND METHODS

### Cell Culture and Treatment

MCF10A mammalian epithelial cells and foreskin fibroblast cells (AG22153, Coriell Institute) were cultured at 37°C in 5% CO_2_ in HuMEC Complete media (Gibco, #12752010) and DMEM media (with 10% FBS and 1% Pen/Strep, VWRL-0101-0500), respectively. Wild-type and p53-knockout HCT116 colon carcinoma and mouse embryonic fibroblasts were kind gifts of Carol Prives and Jing Huang, respectively. To induce p53 activation, cells were treated with Nutlin-3A (5 μM final, dissolved in DMSO) or DMSO (vehicle control) for 6 hours.

### Western Blotting

Cells were lysed in RIPA buffer (10 mM Tris-HCl (pH 8.0), 1 mM EDTA, 1% Triton X-100, 0.1% sodium deoxycholate, 0.1% SDS and 140 mM NaCl, supplemented freshly with protease inhibitors) and probed with antibodies against p53 (BD Biosciences; BD554293), p63 (Cell Signaling; 13109), TA-p63 (BioLegend; 618901), *Δ*Np63 (BioLegend; 619001) and GAPDH (Cell Signaling; 5174).

### RNA-sequencing sample and library preparation

Total mRNA was extracted with E.Z.N.A. Total RNA kit (Omega; R6834-02) and Poly(A)+ RNA was isolated by double selection with poly-dT beads, using 2.5 μg total RNA, which is then followed by first- and second-strand synthesis. Sequencing libraries were prepared using NEXTflex Rapid Illumina DNA-Seq Library Prep Kit (Bioo Scientific). Samples were single-end sequenced on an NextSeq 500. RNA-seq reads were aligned via STAR (24) to Ensembl v.75/hg19. Count tables were generated using STAR-aligned BAM files and HTseq (25). Count tables were then used to call differentially expressed genes using DEseq2 (26).

### Quantitative Real-Time PCR

Total RNA was extracted as per RNA-sequencing protocol and cDNA was synthesized using 1ug of total RNA as a template and the High-Capacity cDNA Reverse Transcription Kit (Applied Biosystems; 413760). Relative standard qPCR was performed using iTaq Universal SYBR Green Supermix (Bio-Rad; 172-5124) with primers shown in Supplementary Table S8 on an ABI 7900HT instrument.

### Lentivirus production, purification and transduction

Lentiviral shRNAs were produced using HEK293T cells that were seeded in 6 well plates. shRNA sequences are as follows: p53 (GTCCAGATGAAGCTCCCAGAA), p63 (CCGTTTCGTCAGAACACACAT). 1 μg of pLKO plasmid having either scramble shRNA or p53 shRNA or p63 shRNA was combined with 1 μg mix of packaging plasmids (pMD2 and psPAX2) and the mixture was diluted into jetPRIME buffer (Polyplus Transfection; 89129-924) and reagents, following the manufacturer’s protocol. Lentivirus containing supernatants were collected at 24 hours and 48 hours post-transfection, and filtered through a 0.45 μm membrane and stored in aliquots at −80°C. MCF10A or SkFib cells were transfected with lentivirus supplemented with 8 μg/ml polybrene. At 24 hour post-infection with lentivirus, media was replaced with the proper puromycin selection (0.5 μg/ml for MCF10A and 2 μg/ml for SkFib).

### ChIP-sequencing sample and library preparation

Cells were crosslinked at 80% confluency with 1% formaldehyde for 10 min at room temperature. Crosslinking was quenched with 125mM glycine and the resulting pellet was washed twice with cold PBS and lysed as previously described (27). Samples were subjected to sonication with Diagenode Bioruptor Plus for 40 cycles (30 sec on/off at high setting) for shearing chromatin to 150-500 bp average size. Immunoprecipitation reactions for SkFib, MCF10A, and HCT116 cell lines were performed with Diagenode IP-Star Compact Automated System, with the exception of p53 ChIP experiments. Mouse embryonic fibroblast and p53 ChIP-seq were performed as described (23). Antibodies were preconjugated to Protein G beads (Invitrogen) against p63 (Cell Signaling; 13109), H3K4me3 (Active Motif; 39159), H3K4me2 (Millipore; 07-030), H3K27ac (Active Motif, 39133) and p53 monoclonal antibody DO1 (BD Biosciences; BD554293). Immunoprecipitated DNA was reverse-crosslinked at 65°C for 4 hours, eluted, purified using SPRI beads, and used to construct sequencing libraries. Sequencing libraries were prepared using NEBNext Ultra DNA Library Prep Kit for Illumina (New England Biolabs). Prior to sequencing, library quality control was performed with Qubit (Thermo Fisher Scientific), Bioanalyzer (Agilent) and qPCR quantification. All ChIP samples, including input, were single-end sequenced on an NextSeq 500 at the University at Albany Center for Functional Genomics. Uniquely aligned reads (up to one mismatch) were aligned to NCBI37/hg19 using Bowtie2 (28).

### ChIP-seq Peak Calling and Differential Binding Analysis

Significant regions of transcription factor (p53, p63) and histone modification (H3K4me2, H3K4me3, H3K27ac) enrichment were called using macs v.2.1.0 (29) preserving only peaks with an adjusted P-value < 0.01. All peak calling was performed with sheared chromatin input. p53 and p63 motif analysis was performed using p53scan (30) and all macs-derived peaks lacking a p53/p63 consensus motif were removed from further analysis. A full list of p53/p63 peaks can be found under GEO Accession Number GSE111009. Global transcription factor motif enrichment experiments were performed using the findMotifsGenome.pl script from HOMER (31). Differential enrichment of p53 ChIP-seq datasets across cell types was performed using DiffBind (32).

### Analysis of histone modification enrichment

Chromatin enrichment analyses were performed using the annotatePeaks.pl script of HOMER. Read-depth-scaled chromatin tag enrichment within a specified window was input-normalized (target – input) and then further scaled to total genomewide enrichment of peaks for that histone modification. H3K27ac ChIP-seq data from MCF7 cells was produced by the ENCODE Project (ENCFF855RCK and ENCFF761EUR). HCT116 and MCF7 p53 ChIP-seq datasets were a part of GSE86164 (3).

### Computational Analysis and Plotting

Peak file intersections were performed using the intersectBed package of BedTools (33), scoring positive intersections as those with at least 1bp of overlap (-f 1E-9). The original peak file (-a) was reported only once (-u) if any intersection exists with the query file (-b). The closestBed package of BedTools was used to measure distances between features with the distance reported using the –d option. Figures were generated using Graphpad Prism or Datagraph. Bigwig (bw) files were generated with the bamCoverage package of deepTools v.2.5.4 (34) with a bin size of 1 (–binSize 1) and read extension (–extendReads 300). Heatmaps were then generated using bigwig files for a -/+ 1000bp region from the p53/p63 peak center using the computeMatrix (reference-point, -a 1000, -b 1000, –binSize 10) and plotHeatmap functions. Genome browser tracks (bedGraph) were generated using the makeUCSCfile.pl package of HOMER (31). K-means clustering was performed using cluster 3.0 for Mac.

### Statistical Testing

Statistical testing was performed using Graphpad Prism (v. 7) or using built-in statistical parameters in *R* (v.3.4.0).

### ChromHMM chromatin state and DNAse hypersensitive site (DHS) enrichment analysis

ChromHMM analysis (35) was performed using the 25 state model (http://egg2.wustl.edu/roadmap/web_portal/), with at least 50% of the p53/p63 motif required to overlap a single chromatin enrichment term (closestBed –a p53.file –b ChromHMM.file –d –f 0.51). The 25 chromatin state models for ChromHMM terms were combined into 4 regulatory terms: enhancer, repressive, quiescent, or transcription. A full list of p53, p63, and p53/p63 binding site regulatory region inferences across the 127 cell types of the ChromHMM analysis can be found in Supplementary Table 10. DHS data were downloaded from http://hgdownload.cse.ucsc.edu/goldenpath/hg19/encodeDCC/wgEncodeRegDnaseClustered/. Intersection of p53/p63 peaks with DHS data was performed using the closetBed package of BedTools, with 100% of the p53/p63 motif required to overlap the DHS (-f 1).

## RESULTS

### p53-dependent transcription and DNA binding are cell type-dependent

We used polyA-enriched RNA-seq to assess gene expression dynamics in model cell lines (MCF10A and SkFib) after p53 activation in response to Nutlin-3A treatment (Figure 1A, Supplementary Figure S1A). We reasoned Nutlin-3A, a non-genotoxic activator of p53, would better reflect p53-dependent transcriptional activity relative to genotoxic activation methods as DNA damage-dependent, but p53-independent, signaling would not confound our analysis (36, 37). Basal and nutlin-3A-induced p53 protein levels are substantially elevated in MCF10A versus SkFib. Consistent with a cell type-dependent activity of p53, MCF10A cells displayed a broad and specific set of Nutlin-3A-induced genes, whereas SkFib showed a markedly smaller, albeit specific, p53-activated transcriptome (Figure 1B). Orthogonal validation by qRT-PCR confirmed a stringent cell type-specificity of each set of genes and demonstrated that canonical p53 targets, like *CDKN1A* and *BTG2* are indeed shared across cell types (Supplementary Figure S1B-D). Decreasing the fold-change cutoff yielded an increased number of differentially regulated genes for each cell type, but maintained the trend that p53-dependent gene targets are more abundant in MCF10A (Supplementary Figure S1E). Genes upregulated in both cell lines are significantly associated with the core p53 response (Supplementary Figure S1F), as expected. MCF10A-specific upregulated genes are enriched in gene ontology groups related to establishment of the skin barrier and p53-dependent processes like programmed cell death and homeostasis (Supplementary Figure S1F). In contrast, SkFib-specific targets are enriched in other cellular stress-related pathways including those associated with the hypoxia response and catabolic processes (Supplementary Figure S1F, Supplementary Table S1). The majority of the MCF10A-specific targets represent novel p53-dependent genes and over 20% of SkFib-specific targets have not been observed previously (Supplementary Figure S1G, Supplementary Tables S2-3 (38)). These initial analyses reveal that the p53-activated transcriptome varies between non-transformed cell types and may reflect tailored, cell type-dependent responses to p53 activation after cellular stress.

**Figure 1.**
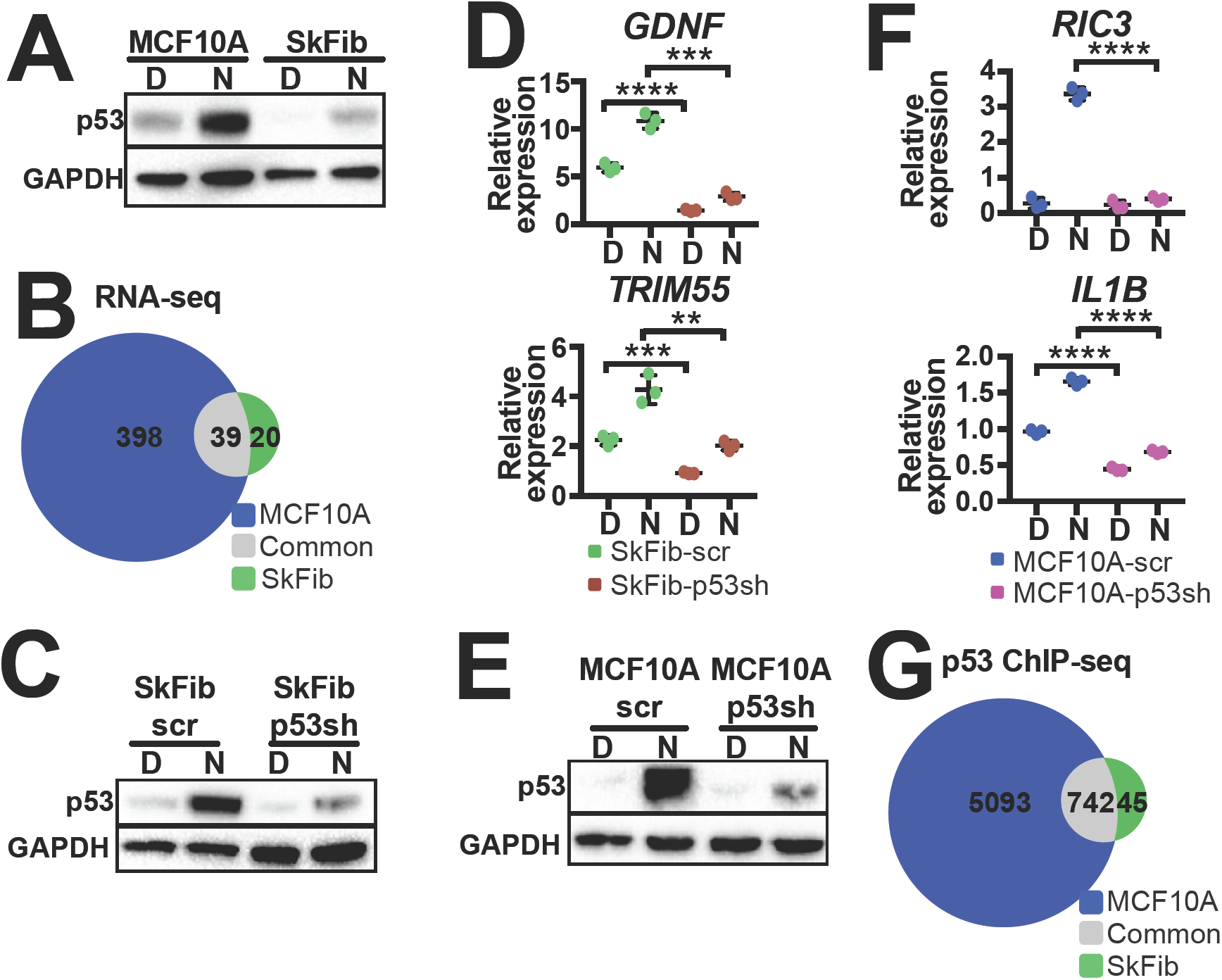
The p53 cistrome and transcriptome are cell type-dependent. (A) Western blot of cell lysates from MCF10A or SkFib cells at 6 hours after DMSO or Nutlin-3A treatment. (B) Venn diagram indicating differentially upregulated genes (Nutlin-3a/DMSO, fold change >2, adjusted p<0.05) in MCF10A or SkFib cells. (C) Immunoblotting for p53 expression at 6 hours of DMSO (D) or Nutlin-3A (N) treatment in SkFib cells stably expressing shRNA against p53 (p53 sh) or a non-targeting control shRNA (scr). (D) qRT-PCR analysis of SkFib-specific gene targets, GDNF and TRIM55, in SkFib cells stably expressing an shRNA to p53 (p53 sh) or a non-targeting control shRNA (scr). Gene expression is normalized to GAPDH expression. Error bars represent SEM; ****p<0.0001, ***p<0.001 and **p<0.01, calculated by Student’s t-test. (E) Immunoblotting for p53 expression at 6 hours of DMSO (D) or Nutlin-3A (N) treatment in response to p53 knockdown in MCF10A cells stably expressing shRNA against p53 or a non-targeting control shRNA (scr). (F) qRT-PCR analysis of MCF10A-specific Nutlin-3A-induced genes, RIC3 and IL1B, in MCF10A cells stably expressing an shRNA to p53 (p53 sh) or a non-targeting control shRNA (scr). Expression is normalized to GAPDH expression. Error bars represent SEM; ****p<0.0001, calculated by Student’s t-test. (G) Venn diagram depicting overlap between significantly-enriched (p<0.01, MACS v2) Nutlin-3A-induced p53 peaks in MCF10A or SkFib.

We next confirmed that select cell type-dependent targets were p53-dependent (Figure 1C-E, Supplementary Figure S1H-I). Indeed, knockdown of p53 in SkFib led to sharp reduction in both basal and Nutlin-3A-induced expression of the SkFib-specific genes, *GDNF* and *TRIM55* (Figure 1D, Supplementary Figure S1H). Depletion of p53 in MCF10A (Figure 1E-F, Supplementary Figure S1I) led to a loss of Nutlin-3A-induced *RIC3* and *IL1B* expression relative to a non-targeting control shRNA (Figure 1F), indicating these genes are *bona fide* p53 target genes in epithelial cells.

Recent analyses reached opposing conclusions with regard to the cell type-dependence of p53 engagement with the genome (21, 39). We thus assessed whether differences in p53 engagement with the genome in our model cell lines might explain our observations of cell type-dependent transcriptomes. Consistent with cell type-dependence, genomewide ChIP-seq revealed a highly-enriched set of MCF10A-specific p53 binding sites and a smaller number of SkFib-specific sites, mirroring our gene expression observations (Figure 1G). We only considered p53 binding sites that replicated across two independent biological replicates. Cell type-dependent differential p53 binding was further validated using the DiffBind statistical framework (Supplementary Figure S1J). p53 enrichment at common sites mirrors both the basal and induced protein expression of p53, with increased enrichment observed in MCF10A relative to SkFib (Figure 1A, Supplementary Figure S1K). p53 protein expression and occupancy differences do not correlate with the ability to activate common targets as SkFib displayed dramatically increased Nutlin-3A induced gene activation compared to MCF10A (Supplementary Figure S1L). Common p53 binding events are primarily within 10kb of genes upregulated in both cell types, similar to previous results suggesting that many direct p53 targets have proximal p53 binding (Supplementary Figure S1M). MCF10A-specific genes have a higher percentage of proximal p53 binding relative to SkFib. Notably, we observe both MCF10A and SkFib p53 binding proximal to SkFib-specific genes (Supplementary Figure S1M). Cell type-specific binding events displayed lower p53 occupancy than at common sites, suggesting that common sites might represent more stable or higher affinity p53 binding sites (Supplementary Figure S1N). This observation is in line with previous results suggesting a core p53 program is well conserved across cell types (3, 21, 40). Taken together, these data provide evidence that the p53-dependent transcriptome is indeed cell type-specific and that p53 engagement with the genome is variable across cell types.

### Promoter and enhancer chromatin states define differential p53 binding and transcriptional activity

We next wanted to identify potential mechanisms driving the strong cell type-specificity within the p53 transcriptome and cistrome. The most straightforward hypothesis is that differential p53 binding drives cell type-specific transcriptome differences, but our data suggest that this might only explain MCF10A behavior (Figure 1G, Supplementary Figure S1M). Thus, we examined *GDNF*, a SkFib-specific gene, and observed proximal p53 binding 8.5kb upstream of the *GDNF* transcriptional start site (Figure 2A). We also noted p53 binding to this region in MCF10A, even though p53-dependent *GDNF* expression is SkFib-specific (Figure 2A). Similar p53 proximal binding was observed at SkFib-specific genes *TRIM55* and *MAMDC2* (Supplementary Figure S2A), suggesting that differential proximal p53 binding *per se* does not explain the transcriptional specificity observed in SkFib relative to MCF10A.

**Figure 2.**
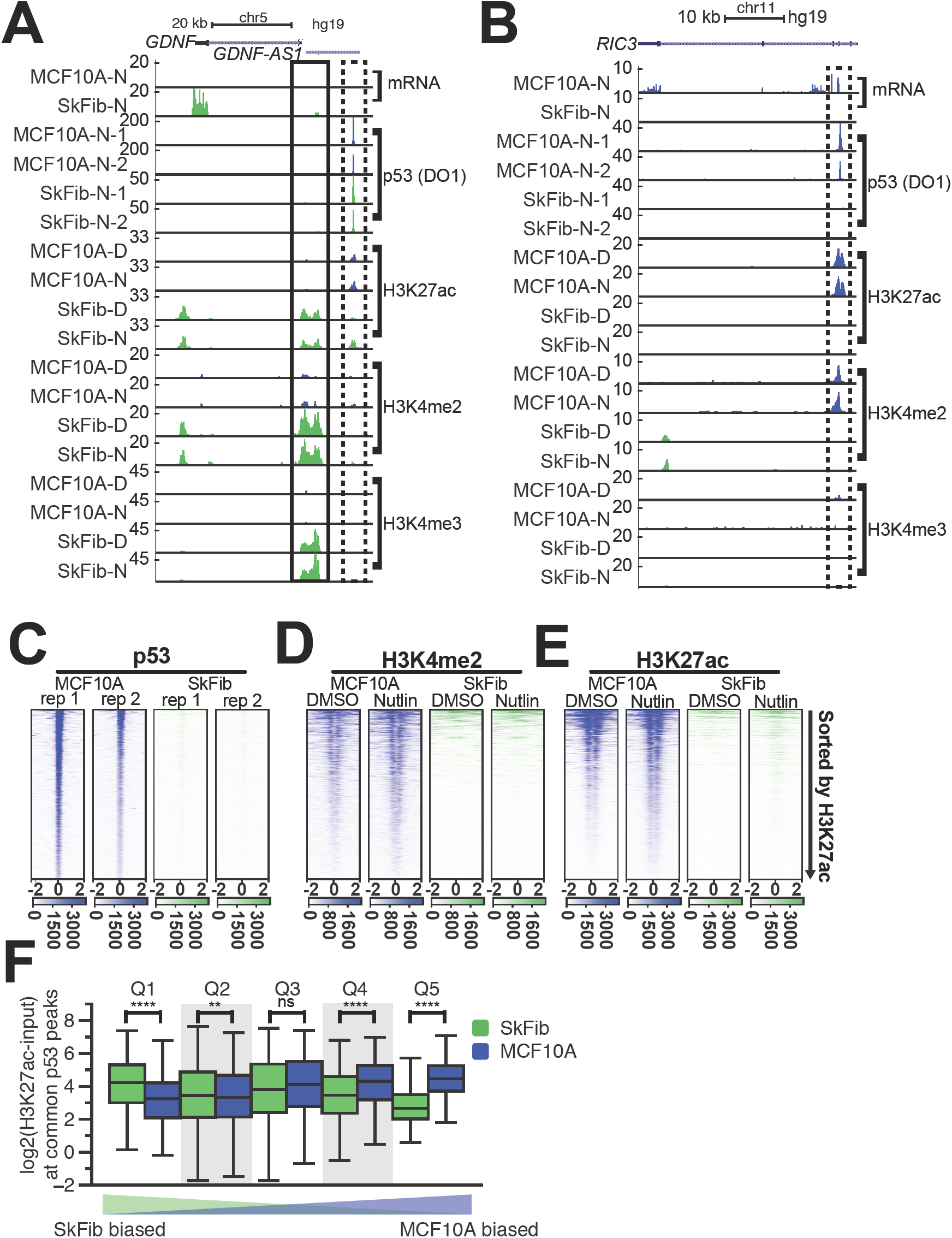
Cis-regulatory activity and differential p53 binding drives cell type-specific target gene expression. (A) Representative UCSC Genome Browser track view of the GDNF locus, a SkFib-specific p53 target. A p53-bound, putative enhancer (H3K27ac+, H3K4me2+, H3K4me3-) is bound by a dashed box, while the putative GDNF promoter (H3K27ac/H3K4me2/H3K4me3+) is bound by a solid box. The y-axis is scaled to the maximum intensity between MCF10A or SkFib for each feature. (B) Representative UCSC Genome Browser track view at the RIC3 locus, illustrating p53 binding to an MCF10A-specific enhancer signature (H3K27ac/H3K4me2+, H3K4me3-) in response to Nutlin-3A treatment. The y-axis is scaled to the maximum intensity between MCF10A or SkFib for each feature. (C) Heatmap plots at MCF10A-specific p53 binding sites for p53 in MCF10A or SkFib within a 4000-bp window (-/+ 2000 bp from the p53 motif; ‘-2’ indicates 2000bp down-stream from the motif, ‘0’ indicates the motif center, and ‘+2’ indicates 2000bp upstream from the motif). Two biological replicates for each dataset are shown. (D) H34me2 and (E) H3K27ac enrichment in DMSO or Nutlin-3A-treated MCF10A or SkFib within a 4000-bp window (-/+ 2000 bp from the p53 motif; ‘-2’ indicates 2000bp downstream from the motif, ‘0’ indicates the motif center, and ‘+2’ indicates 2000bp upstream from the motif). (F) H3K27ac enrichment in a 500bp window from the peak center (input-subtracted, log2) at common rank-ordered SkFib or MCF10A binding sites. ****p<0.0001 and **p<0.01, as calculated by a paired Wilcoxon rank-sum test.

We and others previously proposed that differential enhancer activity may regulate p53-dependent gene expression (23, 41). As enhancers also have a strong cell type-dependence (42, 43), we performed ChIP-seq experiments for canonical promoter (H3K4me3) and enhancer-associated (H3K4me2, H3K27ac) histone modifications to determine whether differentially active promoters or enhancers might partially explain the observed cell type specificity of the p53 transcriptome. The local chromatin environment at the *GDNF* proximal p53 binding site is enriched for H3K27ac and H3K4me2, but lacks H3K4me3, suggestive of an active enhancer in both cell types (dashed box, Figure 2A). Consistent with high basal and p53-inducible expression of *GDNF* in SkFib, the *GDNF* promoter is strongly enriched for the canonical promoter-associated histone modification H3K4me3 (solid box, Figure 2A) which is completely absent from MCF10A. We observed differential promoter/enhancer activity at other SkFib-specific p53 target promoters/enhancers (Supplementary Figure S2A) without differences in p53 binding. Differential promoter/enhancer activity may therefore be a general mechanism by which cell types permit activation of particular p53 target genes.

We next investigated putative mechanisms driving our observation of epithelial-specific p53 target gene activation. p53 binding is pervasive in MCF10A relative to SkFib (Figure 1G), therefore we investigated the connection between *de novo* epithelial p53 binding and the induction of specific gene targets. Examination of the MCF10A-specific targets *RIC3* and *MYBPHL* revealed MCF10A-specific binding of p53 to putative intragenic enhancers (Figure 2B, Supplementary Figure S2B). These putative enhancer regions, characterized by enrichment of H3K4me2 and H3K27ac, and depleted H3K4me3, appeared specific to MCF10A cells. Globally, MCF10A-specific p53 binding occurs within regions that are strongly enriched for this chromatin modification-based signature of transcriptional enhancers (Figure 2C-E) whereas these same genomic regions lack p53 binding and chromatin features in SkFib (Figure 2C-DE, Supplementary Figure 3SB-E).

Our results suggest that p53 binds to genomic loci enriched with enhancer-associated chromatin modifications in MCF10A, but the mechanisms leading to this observation are not known. Almost all shared p53 binding sites (708/720) display higher ChIP-seq signal in MCF10A relative to SkFib (median=5.3X enrichment, Supplementary Figure S1K). Increased p53 protein expression levels presumably drive higher p53 occupancy at these common binding events in MCF10A (Figure 1A and Supplemental Figure S1M, common), but whether the increased expression drives p53 binding affinity/dynamics across cell types is unknown. Because of the uniformly high enrichment in MCF10A relative to SkFib, we rank ordered shared p53 binding site enrichment values within cell types and then compared ranks across cell types. p53 enrichment ranks were well correlated between biological replicates (0.961 for MCF10A and 0.852 for SkFib, Spearman rho). Rank-orders were less well-correlated when comparing across cell types (MCF10A vs. SkFib, 0.628, Spearman rho) with notable outliers (Supplementary Figure S3F). p53 occupancy at an intragenic region of *ARID3A* is similar in both MCF10A and SkFib despite significantly higher overall occupancy in MCF10A for the majority of binding sites (Supplementary Figure S3G). Local H3K27ac and H3K4me2 are also more highly enriched at this loci in SkFib relative to MCF10A, leading us to ask whether local chromatin structure influences the relative occupancy of p53 binding sites.

To further address this possibility, we grouped common p53 binding events into quintiles based on differences in the rank-order of enrichment scores between MCF10A and SkFib and then assessed local chromatin features. Common p53 binding sites with SkFib-biased binding displayed significantly more enrichment of H3K27ac and H3K4me2 in SkFib than in MCF10A (Figure 2F, Supplementary Figure 3H). Similarly, MCF10A-biased p53 binding locations have significantly more H3K27ac and H3K4me2 enrichment in MCF10A. p53 binding sites lacking cell type-bias showed no enhancer-associated chromatin enrichment differences (Figure 2F). Comparison of biased p53 binding and H3K27ac enrichment in MCF7 and HCT116 cell lines yielded similar chromatin enrichment bias (Supplementary Figure 3I). Overall, these observations implicate local chromatin structure state, specifically enhancer-associated histone modifications, in p53 binding site selection within and across cell types.

Enhancer specification is generally thought to be regulated by the combined effort of both common and lineage-dependent transcription factors (43). Multiple recent reports suggest that p53 activity at *cis-*regulatory elements is independent of any accessory transcription factors, primarily due to low enrichment/specificity of other DNA motifs (21, 22, 41). Indeed, MCF10A enhancers are highly enriched for underlying p53/p63 motifs relative to SkFib, consistent with results from the FANTOM consortium and a lack of any obvious epithelial-specific transcription factor motifs (Supplementary Figure S3J, Supplementary Table S4, (44)). As the p53 family shares a common, although not identical, response element motif, we asked whether p53 family members might regulate MCF10A-specific p53 binding events, enhancer-associated chromatin modifications, and gene expression.

### p53 and p63 cooperate at enhancers to regulate epithelial-specific gene expression

Epithelial cells highly express p63 and are dependent on p63 activity for self-renewal and epidermal commitment *in vivo* (45). p63 also bookmarks enhancers during epidermal differentiation (46) and may regulate chromatin accessibility through interactions with chromatin modifiers (47, 48). ΔN-p63, not TA-p63, is expressed in MCF10A which also have undetectable levels of the other p53 family member p73 (Supplementary Figure S4A-D). Therefore, we focused on the function of p63 in epithelial enhancer activity by performing genomewide ChIP-seq of p63*α* in MCF10A after DMSO or Nutlin-3A treatment. Examination of *IL1B* and *RNASE7*, MCF10A-specific p53 targets, displayed proximal p63 binding to distinct enhancer regions co-occupied by p53 in an MCF10A-specific manner (Figure 3A, Supplementary Figure S4E). Globally, p63 binds to significantly more genomic loci than does p53 (Figure 3B), and over 75% of epithelial-specific p53 binding events overlap p63 binding. Shared binding sites display higher occupancy and are more likely to have enhancer-associated chromatin modifications than do sites unique to either p53 or p63 (Figure 3C, Supplementary Figure S4F), suggesting that both p53 and p63 might be necessary for full enhancer activity.

**Figure 3.**
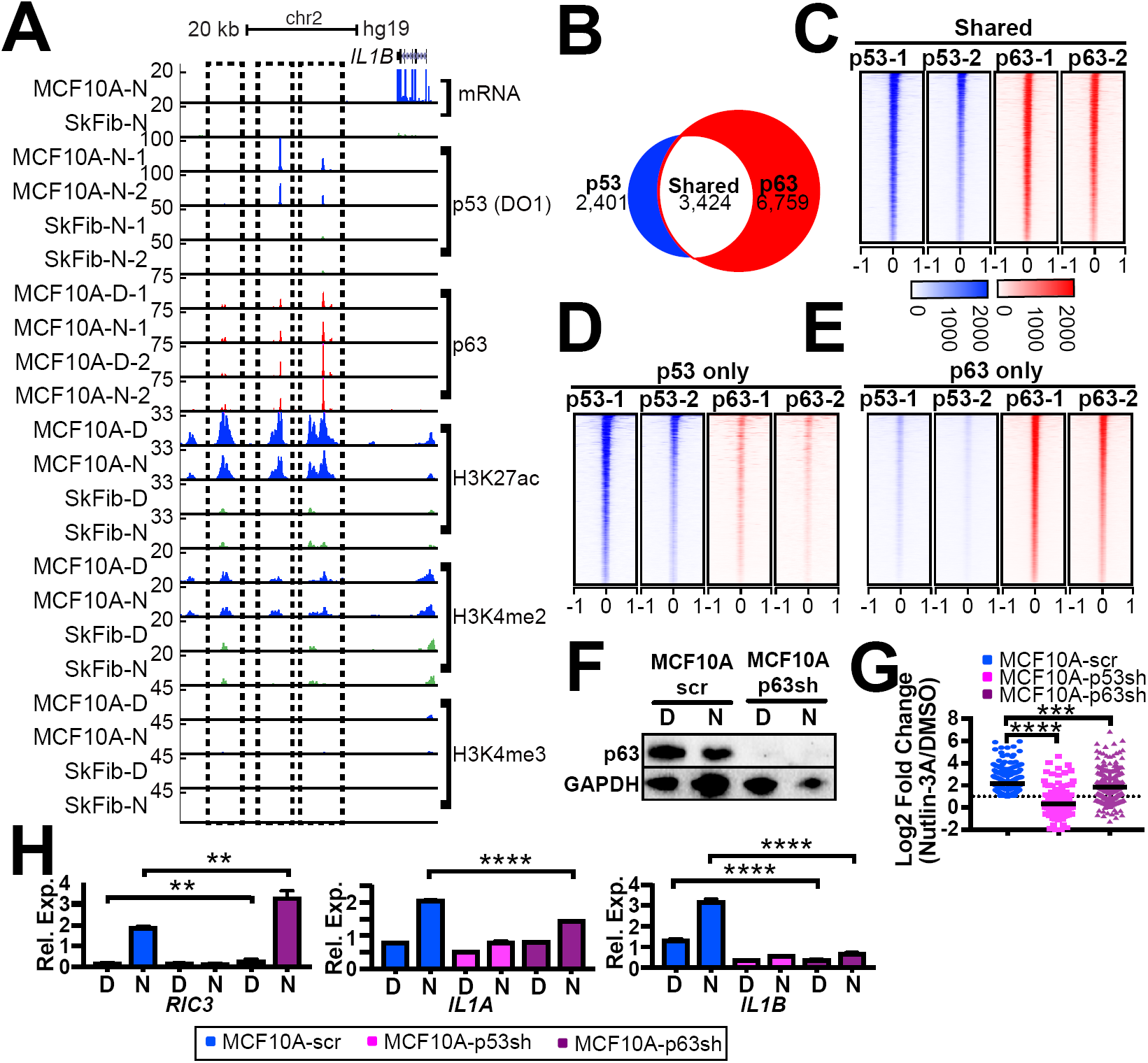
p63 engages with p53-bound enhancers and regulates p53-dependent transactivation of epithelial target genes. (A) Representative UCSC Genome Browser track view of the IL1B locus, illustrating three MCF10A-specific putative enhancers bound by p53 and p63 in response to Nutlin-3A treatment (H3K27ac+, H3K4me2+, H3K4me3-; dashed box). (B) Venn diagram representation of the overlap between biological replicates of p53 and p63 ChIP-seq peaks (input-normalized, p53/p63 motif-positive, MACS v2, p<0.01) in MCF10A when treated with Nutlin-3A. Heatmap plots of p53 and p63 enrichment at (C) shared, (D) p53 only, and (E) p63 only binding sites in MCF10A cells within a 2000-bp window (-/+ 1000 bp from the peak center) in response to Nutlin-3A treatment. (F) Immunoblotting for p63 in response to p63 knockdown in MCF10A cells stably expressing shRNA to p63 or a non-targeting control shRNA (scr) after 6 hours of DMSO (D) or Nutlin-3A (N) treatment. (G) Fold-change (Nutlin-3A/DMSO, log2, median in black) of MCF10A-specific Nutlin-3A-induced genes in MCF10A cells expressing shRNA targeting either a non-targeting control (scr), p53, or p63. ****p<0.0001 and ***p<0.001 (H) qRT-PCR analysis of MCF10A-specific Nutlin-3A-induced genes, RIC3, IL1A and IL1B. Expression is normalized to GAPDH expression. Error bars represent SEM; ****p<0.0001 and **p<0.01, calculated by Student’s t-test.

Overall, p63 binding correlates with epithelial-specific enhancer modifications and co-occupies the majority of p53 sites (Figure 3B-E, Supplementary Figure S4G). The modal binding of p63 is within 10kB of the TSS of Nutlin-3A-induced genes, further suggesting that p63 might co-regulate these specific target genes (Supplementary Figure S4G). In order to test whether p63 regulates p53 activity, we created MCF10A cells stably-expressing shRNA against p63 (Figure 3F, Supplementary Figure S4H). We then performed RNA-seq analysis of MCF10A cells expressing non-targeting (scr), p53, or p63 shRNA to determine their relative contributions to epithelial-specific target genes. Loss of p53 expression strongly inhibited Nutlin-3A-induced gene expression consistent with this set of genes representing true p53 targets (Figure 3G). Knockdown of p63 showed more pleiotropic effects. Globally, Nutlin-3A-induced target gene activation is inhibited in p63 knockdown relative to non-targeting controls, suggesting p63 activity is required for a set of epithelial p53 targets (Figure 3G). Whereas 40% of p53 target genes show defective induction in the absence of p63, the remainder of the genes are either unaffected or display increased p53-dependent transactivation (Supplementary Figure S4I). This observation is consistent with previous reports of direct repression of p53-dependent transactivation through binding site competition (18, 19).

Our data suggest that p63 also functions to co-activate a set of p53-dependent genes specifically in epithelial cell types. We then validated these results by qRT-PCR on a panel of epithelial target genes after p53 or p63 knockdown. *RIC3,* previously demonstrated to be a p53-dependent gene (Figure 1F), is also derepressed and shows higher p53-dependent transactivation in the absence of p63 (Figure 3H). Conversely, both *IL1A* and *IL1B* are dependent on p53 and p63 for full transactivation, with *IL1B* showing a strong dependence on p63 for basal expression (Figure 3H). Knockdown of p63 also led to basal gene expression changes for p53 targets (Supplementary Figure S4J) and genes related to cell cycle and cell adhesion/extracellular matrix (Supplementary Table S5), as has previously been described (49). In total, these results suggest that p63 functions as both a co-activator and repressor of p53-dependent transcription in MCF10A, but how these opposing mechanisms lead to differential p53 activity is not clear.

### Varying roles for p53 family members in regulation of enhancer identity

p53 and p63 engagement to the genome in epithelial cells correlates with enhancer activity and the p53 family are putative pioneer factors with enhancer bookmarking activity (23, 46, 47, 50). Therefore, we examined the effect of p53 and p63 on enhancer identity in MCF10A. Globally, loss of p63 led to a reduction of H3K27ac and H3K4me2 at p63-bound enhancers (Figure 4A-B). Global H3K4me2 enrichment at enhancers was unaffected by p63 loss (Figure 4B) while H3K27ac levels at enhancers were generally increased (Figure 4A). Expression of *RRM1*, a member of the ribonucleotide reductase enzyme complex, was significantly downregulated by loss of p63 and unaffected in the absence of p53 (Figure 4C). p63-bound enhancers upstream of *RRM1* and *EDN2*, another epithelial p53 target gene, showed dramatic loss of H3K27ac and H3K4me2 enrichment in p63-depleted cells, with similar loss of enhancer-associated modifications observed broadly at p63 binding sites (Figure 4A-C, Supplementary Figure S5A). In total, over 20% of H3K4me2+ and 15% of H3K27ac+ p63-bound sites showed at least a 2-fold depletion in MCF10A-p63sh cells relative to scrambled shRNA (Figure 4D). These enhancer-associated histone modifications were generally unaffected by the loss of p53 (Figure 4D), suggesting this behavior is specific to p63. This loss of enhancer-associated histone modifications at p63 binding sites suggests that p63 plays a direct role in the maintenance of enhancer activity.

**Figure 4.**
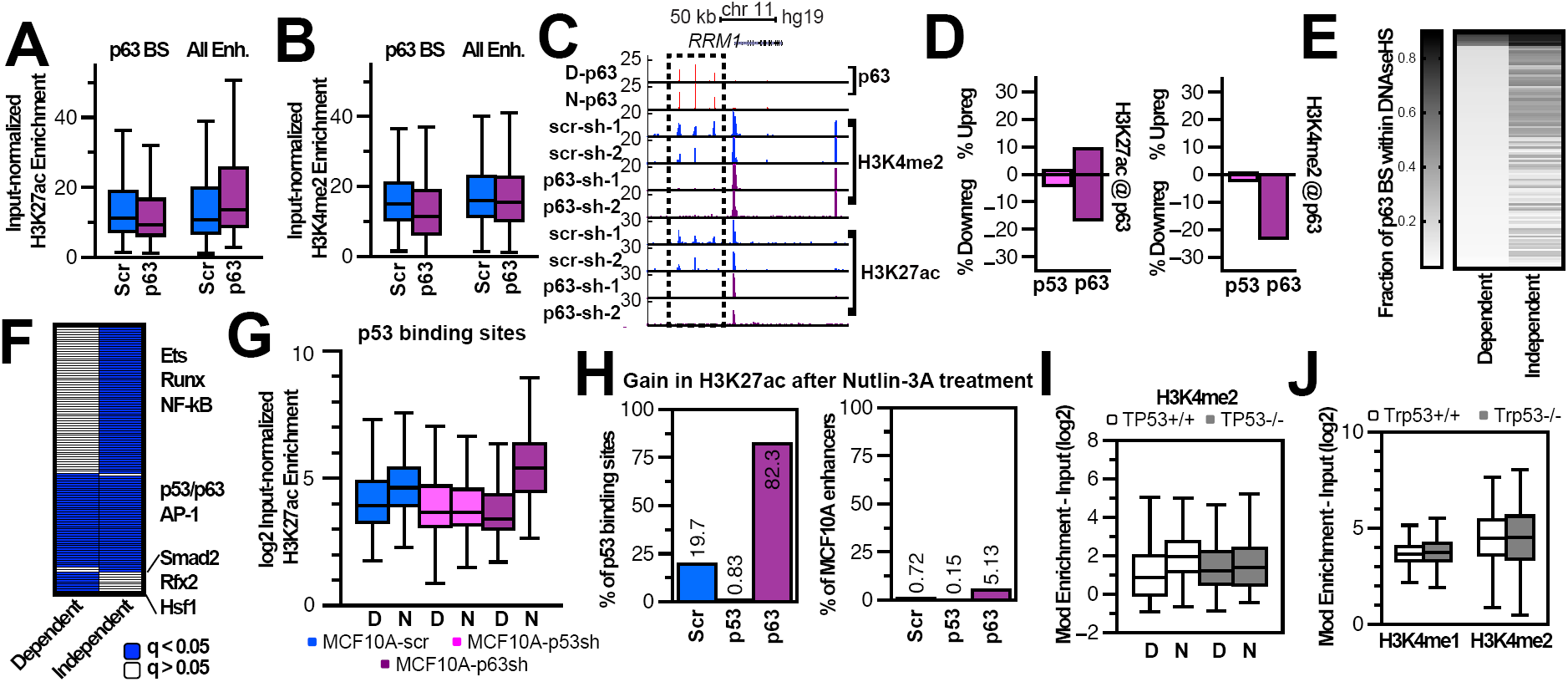
p63, and not p53, maintains enhancer chromatin identity. Input-subtracted (A) H327ac or (B) H3K4me2 enrichment at p63 binding sites (left, -/+ 250bp from p63 peak center) or at all remaining enhancers (H3K27ac+, H3K4me2+, H3K4me3-, right) in MCF10A cells expressing non-targeting (Scr) or p63 shRNA. (C) Representative UCSC Genome Browser track view of the RRM1 locus, illustrating three MCF10A-specific putative enhancers bound by p63 that are lost in response to p63 depletion (H3K27ac+, H3K4me2+, H3K4me3-; dashed box). (D) Bar graphs depicting the percent of p63 peaks that show 2-fold gains or losses of H3K27ac (left) or H3K4me2 (right) in response to either p53 or p63 depletion relative to non-targeting control shRNA. (E) Heat-map depicting the fraction of p63 binding sites that lose H3K4me2 (left) or remain constant (right) after p63 depletion found within regions of DNAse hypersensitivity (DHS) across multiple cell lines. (F) Heatmap of k-means clustered q-values for motif enrichment found at p63-dependent or independent enhancers. (G) H3K27ac enrichment at p53 binding sites in MCF10A cells expressing the represented shRNA molecules after 6hours of DMSO (D) or 5uM Nutlin-3A (N) treatment. (H) Bar graph displaying the number of p53 binding sites (left) or total cellular complement of H3K27ac+/ H3K4me2+/H3K4me3-enhancers with more than 2-fold change in H3K27ac enrichment after Nutlin-3A treatment of MCF10 cells expressing the indicated shRNA. (I) H3K4me2 enrichment at p53 binding sites (20) in HCT116 TP53+/+ or -/- cells in response to DMSO or Nutlin-3A treatment. (J) H3K4me1 and H3K4me2 enrichment at p53 binding sites (67) in Trp53+/+ or Trp53-/- mouse embryonic fibroblasts.

Our observations suggest that p63 activity is required for H3K27ac and H3K4me2 enrichment at a subset of its bound enhancers. p63-bound enhancers dependent on p63 for H3K4me2 enrichment (p63-dependent) were significantly more likely to overlap epithelial-specific DNase hypersensitive sites than those bound by p63, but not affected by p63 depletion (p63-independent, Supplementary Figure S4B, Supplementary Table S6). p63-dependent enhancers are DNAse-hypersensitive specifically in epithelial cells whereas p63-independent enhancers appear to be more broadly accessible across cell types (Figure 4E). Overall, p63 binding sites are strongly enriched for enhancer-associated modifications across epithelial cell types consistent with its well-established expression pattern and role as a lineage determination and pioneer transcription factor (Supplementary Figure S4C).

Enhancer accessibility and activity is controlled by combinatorial transcription factor binding (51, 52), thus we compared enrichment of transcription factor motifs across p63-dependent and independent enhancers. Both types of p63-bound enhancers are enriched for p53/p63 and AP-1 family motifs (Figure 4F), and p63- dependent enhancers display limited enriched motifs like the SMAD family. Conversely, p63-independent enhancers are strongly enriched for Ets family motifs relative to p63-dependent enhancers (Figure 4F, Supplementary Table S7). These data are consistent with the broad chromatin accessibility of p63-independent enhancers observed across cell types. Therefore, our results demonstrate that p63 activity is critical for local H3K27ac and H3K4me2 enrichment at a set of epithelial-specific enhancers and that p63 binds to another set of cell type-independent enhancers putatively dependent on other transcription factor families like ETS that the activity of p63-independent enhancers is dependent on other transcription factors. Combinatorial transcription factor binding at enhancers is critical for appropriate spatial and temporal gene expression (51, 52), and the collaborators at p53 family-bound enhancers are relatively understudied compared to promoters (41). Recent *in vivo* dissection of p53 enhancer activity implicated the general enhancer-associated AP-1 family member CEBPB as a direct regulator of an enhancer upstream of *CDKN1A* (53), thus contributions of other transcription factors to p53 family enhancer function are of particular interest.

Our data indicates that p63 likely functions to maintain chromatin structure and activity at a set of epithelial-specific enhancers, but not for the entire complement of p63-bound enhancers. A number of studies have demonstrated that p53 has pioneer factor activity as it can directly bind its consensus sequence in the context of a nucleosome (23, 54, 55). p53 may possess pioneer factor activity, but the large majority of p53 binding sites are constitutively closed or are within previously accessible chromatin (3, 22, 23). Therefore, we examined whether p53 might serve pioneer or maintenance roles at enhancers as well. Basal H3K4me2 and H3K27ac enrichment at p53 binding sites was unaffected by the loss of p53 (Fig 4G, Supplementary Figure 5D). p63 knockdown led to a slight loss of H3K27ac and H3K4me2 enrichment at p53 binding sites (Figure 4G, Supplementary Figure S4D), an expected result as approximately 75% of p53 binding events in MCF10A cells are also p63 binding sites (Figure 3B). Examination of p53+/p63- binding events displayed a similar trend, with both H3K27ac and H3K4me2 unaffected by p53 loss (Supplementary Figure S4E). These results suggest that p53 plays a limited role in the specification and maintenance of basal enhancer. The observed p53-independence of basal enhancers is consistent with previous observations that p53 is dispensable for basal eRNA abundance at enhancers (37). Unlike its role in basal enhancer activity, p53 is absolutely required for *de novo* Nutlin-3A-induced H3K27ac levels at p53-bound enhancers (Figure 4G-H). Depletion of p63 results in a substantial increase in Nutlin-3A-induced H3K27ac at p53 binding sites (Figure 4G). Nearly 85% of p53-bound enhancers gain H3K27ac (> 2-fold relative to DMSO) in the absence of p63, suggesting that p63 binding can locally inhibit p53-dependent chromatin modification enrichment and activity.

Epithelial cell types like MCF10A express multiple p53 family members that might redundantly regulate enhancer activity, making it difficult to assess the direct role of p53 in modulating enhancer activity. We therefore extended our analysis to isogenic WT or *TP53*-/- HCT116 human colon carcinoma cell lines and WT or *Trp53*-/- mouse embryonic fibroblasts (MEF). Both cell models lack endogenous p63 expression and thus should allow us to define the direct role of p53 in enhancer maintenance and activity. Similar to our observations in MCF10A cells, basal enhancer-associated H3K4me1/2 methylation at p53-bound enhancers is unaffected by the loss of p53 (Figure 4I-J). These data suggest that p53 is not an enhancer maintenance factor and contributes primarily to *de novo* histone acetylation and transcriptional activation at already established enhancers.

## DISCUSSION

In this report, we focused on identifying mechanisms that control differential expression of p53 target genes across cell types. Our results demonstrate that differential *cis-*regulatory activity and p53 binding serve as critical determinants of cell type-specific, p53-dependent transcriptomes. The acute p53 response in foreskin fibroblasts and breast epithelial cells (MCF10A) leads to distinct transcriptional programs that appear to reflect cell type-specific functions. In the case of MCF10A, p53 activates a series of genes involved directly in epithelial cell identity, such as epithelial cornification genes and ZNF750, a key regulator of epithelial cell differentiation (56, 57). Knockout studies in mice suggest p53’s role in development is limited, although neural tube and germ cell defects are observed at low penetrance (58–60). Therefore, our observations that p53 regulates key epithelial development and differentiation genes is surprising. Loss of p53 activity is a strong predictor of epithelial-to-mesenchymal transition (EMT) (61, 62). One potential rationale for p53-dependent regulation of these key epithelial lineage genes is a mechanism to protect against EMT by supporting transcription of epithelial identity genes. Of note, canonical p53 transcriptional pathways appear to be dispensable for tumor suppression (4). Recent work demonstrated that the p53 transcriptional network is highly distributed as to allow loss of multiple target genes/pathways without loss of p53-dependent tumor suppression (3). Certainly, additional investigations into cell type-dependent p53 transcriptional targets, and their roles in context-dependent tumor suppression are warranted given these findings.

Our data suggest that transcriptional activation by p53 requires both gene-proximal p53 binding and pre-establishment of *cis-*regulatory elements. Fibroblast-specific p53 target genes have active promoters, characterized by canonical H3K4me3 enrichment, and p53 engagement with distal enhancer regions. Conversely, the same enhancers were active and engaged by p53 in MCF10A, but the gene promoters had chromatin-associated hallmarks of inactivity. It is currently unclear whether the lack of activity in MCF10A is driven by lack of promoter-bound cofactors, through repressive chromatin-associated mechanisms, or both. Repressive DNA and histone modifications have been implicated in differential control of p53 activity in cancer cell lines (63–65). These observations suggest that promoter competence, independent of p53 binding, potently controls cell specific p53 transcription responses.

Although p53 is broadly expressed in all cells, distal, p53-bound enhancers display significant differences in chromatin structure across cell types (23). Our results suggest that p53 is not required for establishment or maintenance of chromatin modification patterns at these enhancers. p53 is a pioneer transcription factor capable of binding its response element within nucleosomal DNA (50, 54). Indeed our results demonstrate that the majority of p53 binding events occur within nucleosome rich regions lacking active histone modifications in agreement with previous work (23). We propose that p53, which can bind to closed chromatin, does not broadly control *de novo* enhancer accessibility. A recent ATAC-seq study in primary human fibroblasts demonstrated increased chromatin accessibility at a limited number of p53 binding sites (22). Taken together, these data and our observations suggest that p53’s role in mediating enhancer accessibility may be context-dependent and depend on the presence of appropriate chromatin remodelers and other cofactors. Our data also support a model in which the combination of locally-bound cofactors and chromatin environment influence p53 engagement with the genome. The large disparity in the number of p53 binding sites between MCF10A breast epithelial cells and dermal fibroblasts correlates with cell type-dependent differences in local chromatin modification patterns. While p53 protein is more abundant in MCF10A, basal enhancer-associated chromatin modification states (ie those before p53 binding) do not appear to be influenced by the presence of absence of p53. Additionally, the occupancy/strength of p53 binding events found in both cell types is biased towards sites with higher enrichment of enhancer-associated chromatin modifications. These observations suggest that p53, although capable of binding to inaccessible chromatin, is strongly influenced by local chromatin accessibility and enhancer activity.

Initial observations of p53:nucleosomal DNA binding actually demonstrated higher levels of p53-dependent transcriptional activation than when bound to naked/accessible DNA (55). This pioneer factor activity was confirmed in numerous genomewide and *in vitro* biochemical studies. Of note, not all nucleosome rotational positions permit p53 engagement with its response element which may represent a method of repressing p53 activity at the level of chromatin (50). p63 appears to share similar nucleosomal binding activities as p53, although detailed biochemical experiments have not yet been undertaken (66, 67). Indeed, our data suggests that p63 binds to many locations that appear to be nucleosome rich, although whether p63 is directly engaging nucleosomal DNA is unclear. The context-specific nucleosomal DNA binding activity of the p53 family, therefore, may be important for both facilitating robust transcriptional activation and for locally regulating p53 family transcription factor activity at chromatin. These data support a model whereby the p53-dependent transcriptome is licensed by the enhancer regulatory abilities of other transcription factors. This model is a particularly attractive avenue for further inquiry and provides a straightforward mechanism for cell type-dependent tumor suppressor and homeostatic activities of p53.

p53 family members have highly conserved DNA binding domains that allow them to engage with highly similar DNA sequences. Therefore, the extent to which p53 family member competition for DNA binding affects their function has been a longstanding question. Initial observations suggested that p63 primarily represses p53 activity in a dominant negative fashion (18, 19). We observed p63-dependent repression of nearly 15% of p53 targets in MCF10A epithelial cells, supporting a dominant negative model. However, our results also demonstrated that p63 is required for p53-dependent activation of approximately 40% of epithelial target genes. The precise mechanisms by which p63 supports p53 activity are unclear, but our data demonstrating that p63 likely functions as an enhancer licensing or maintenance factor provides a straightforward possibility. Our model proposes that p63 first establishes a permissive chromatin environment which then allows additional transcription factors, including p53, to bind. This putative mechanism supports both p63-dependent p53 activity and p63-mediated repression. Whereas p63 activity may be required to facilitate a suitable binding environment, ΔNp63 could also repress p53 through local competition for the same binding site. We note that loss of p63 led to substantial gains in p53- dependent H3K27ac enrichment at enhancers, supporting a local inhibition of p53 enhancer activity by p63. Due to the population-based nature of ChIP-seq, it is uncertain whether p53 and p63 truly compete for binding sites in the same cell, or whether p53 and p63 binding occurs in mutually exclusive cells populations. Therefore, our data confirm p53 and p63 have their previously reported antagonistic relationship and reveal a more broadly cooperative partnership than expected (8, 9).

Our data demonstrate that p63 is required for enhancer identity in breast basal epithelial cells. Knockdown of p63 in MCF10A cells leads to a site-specific loss of enhancer-associated histone modifications and linked gene expression. p63 is required for lineage commitment and self-renewal of basal epithelial cells (45), and p53 family motifs are highly and specifically enriched in epithelial specific enhancers (44). The specific molecular events linking initial p63 DNA binding to establishment of an active enhancer state are becoming more clear. p63 interacts with the histone H3K4 monomethylase KMT2D in normal epithelial keratinocytes and reduced KMT2D activity leads to preferential reduction of H3K4me1 at p63-bound enhancers (48). Similarly, inactivation of the BAF chromatin remodeling complex is correlated with reduced chromatin accessibility at p63 binding sites (47). Our data, therefore, place p63 in line with other lineage determination factors that directly act as pioneer transcription factors to license cell type-specific enhancers (68, 69). Further investigation into p63 control of epithelial enhancer identity is required to determine precisely how p63 pioneer activity controls epithelial lineage determination and self-renewal.

## ACCESSION NUMBERS

All raw data associated with this study are deposited at NCBI Gene Expression Omnibus under accession number GSE111009.

## ACKNOWLEDGEMENTS

We are grateful to Allison Catizone and Enrique Lin Shiao for feedback on this manuscript. We thank Jason Herschkowitz, Carol Prives, and Jing Huang for the MCF10A, HCT116 p53 KO, and MEF p53 KO cell lines, respectively. Finally, we acknowledge the University at Albany Center for Functional Genomics for sequencing support.

---

**Supplemental Figure S1.**
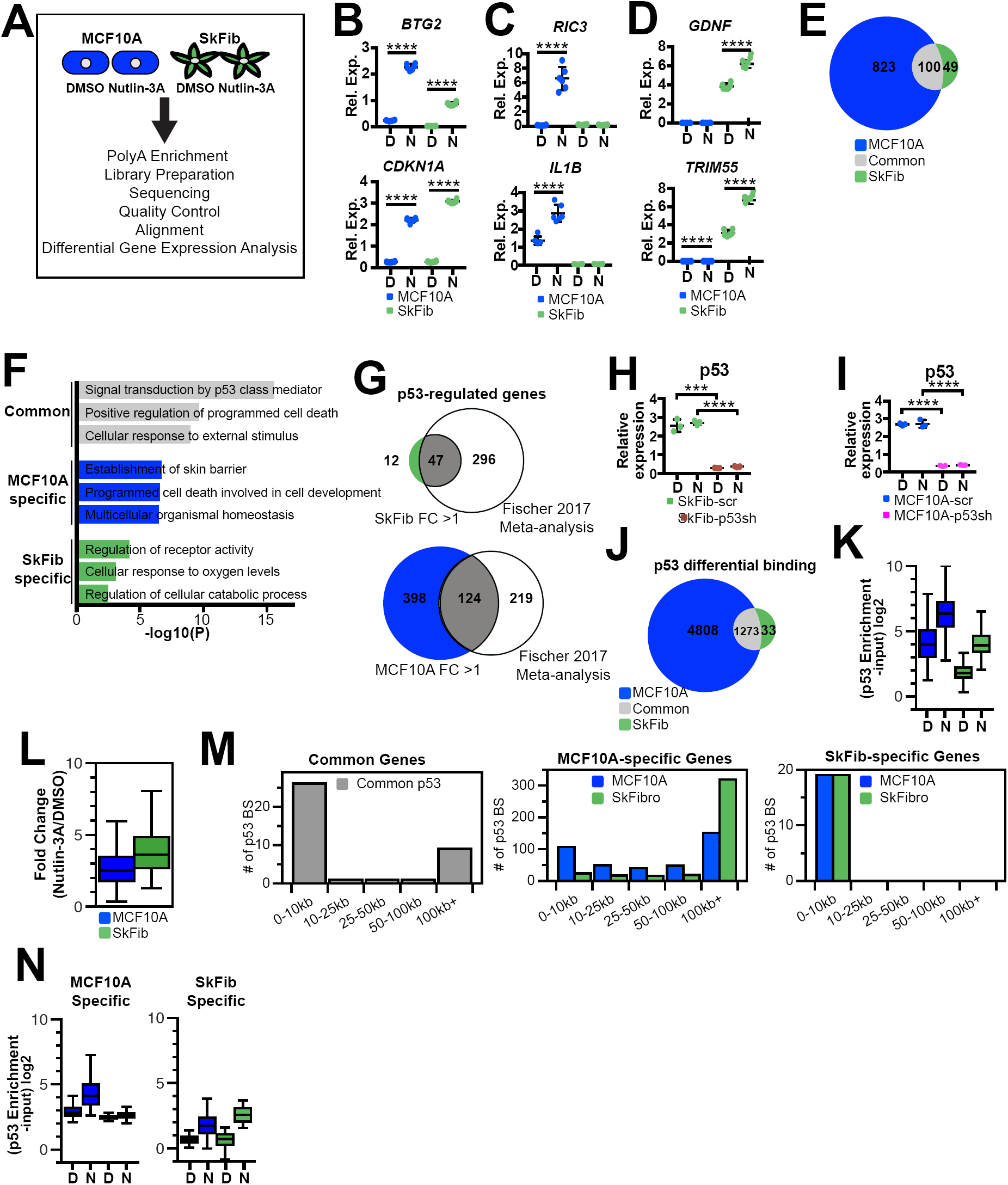
Cell type-dependent p53 transcription in MCF10A and SkFib cells. (A) Experimental design for RNA-sequencing analysis in response to p53 activation by DMSO or 5µM Nutlin-3A for 6 hours. (B-D) qRT-PCR validation of two representative (B) common, (C) MCF10A-, and (D) SkFib-specific Nutlin-3A-induced targets. Target gene expression is normalized to GAPDH and data represent technical replicates from three independent biological replicates. Error bars represent SEM with p values calculated by Student’s t-test, ****p<0.0001. (E) Venn diagram indicating differentially upregulated genes (Nutlin-3a/DMSO, fold change □2, adjusted p<0.05) in MCF10A or SkFib cells. (F) Gene ontology analysis of differentially expressed genes from MCF10A and SkFib cells showing upregulated genes for top three significant biological processes. (G) Intersection between SkFib (top) or MCF10A (bottom) p53 targets identified in this work compared to the meta-analysis of p53 targets identified in at least 3 independent experiments (Fischer, M 2017). (H) qRT-PCR analysis of p53 expression at 6 hours of DMSO (D) or Nutlin-3A (N) treatment in SkFib cells stably expressing shRNA against p53 (p53 sh) or a non-targeting control shRNA (scr). p53 expression is normalized to GAPDH expression (Relative expression). Error bars represent SEM; ****p<0.0001 and ***p<0.001, calculated by Student’s t-test. (I) qRT-PCR analysis of p53 expression at 6 hours of DMSO (D) or Nutlin-3A (N) treatment in response to p53 knockdown in MCF10A cells stably expressing shRNA against p53 or a non-targeting control shRNA (scr). p53 RNA expression is normalized to GAPDH expression. Error bars represent SEM; ****p<0.0001, calculated by Student’s t-test. (J) Venn diagram depicting differentially bound p53 peaks in SkFib and MCF10A according to replicate analysis using DiffBind. (K) Boxplots depicting enrichment of p53 (input-normalized, log2) at common p53 binding sites found in both MCF10A (blue, left) and SkFib (green, right). (L) Fold change ratio of Nutlin-3A/DMSO of common input-subtracted p53 enrichment for MCF10A or SkFib. (M) The percentage of p53 peaks common in MCF10A and SkFib (left), or only in MCF10A (middle), or only in SkFib (right) in response to Nutlin-3A treatment within varying distances to the nearest transcription start site (TSS) of a Nutlin-3A-induced gene. (N) Boxplot analysis of the input-subtracted p53 enrichment for MCF10A (blue, left) or SkFib (green, right) for MCF10A-specific-(left) or SkFib-specific-p53 binding sites (right) in response to DMSO (D) or Nutlin-3A (N) treatment.

**Supplemental Figure S2.**
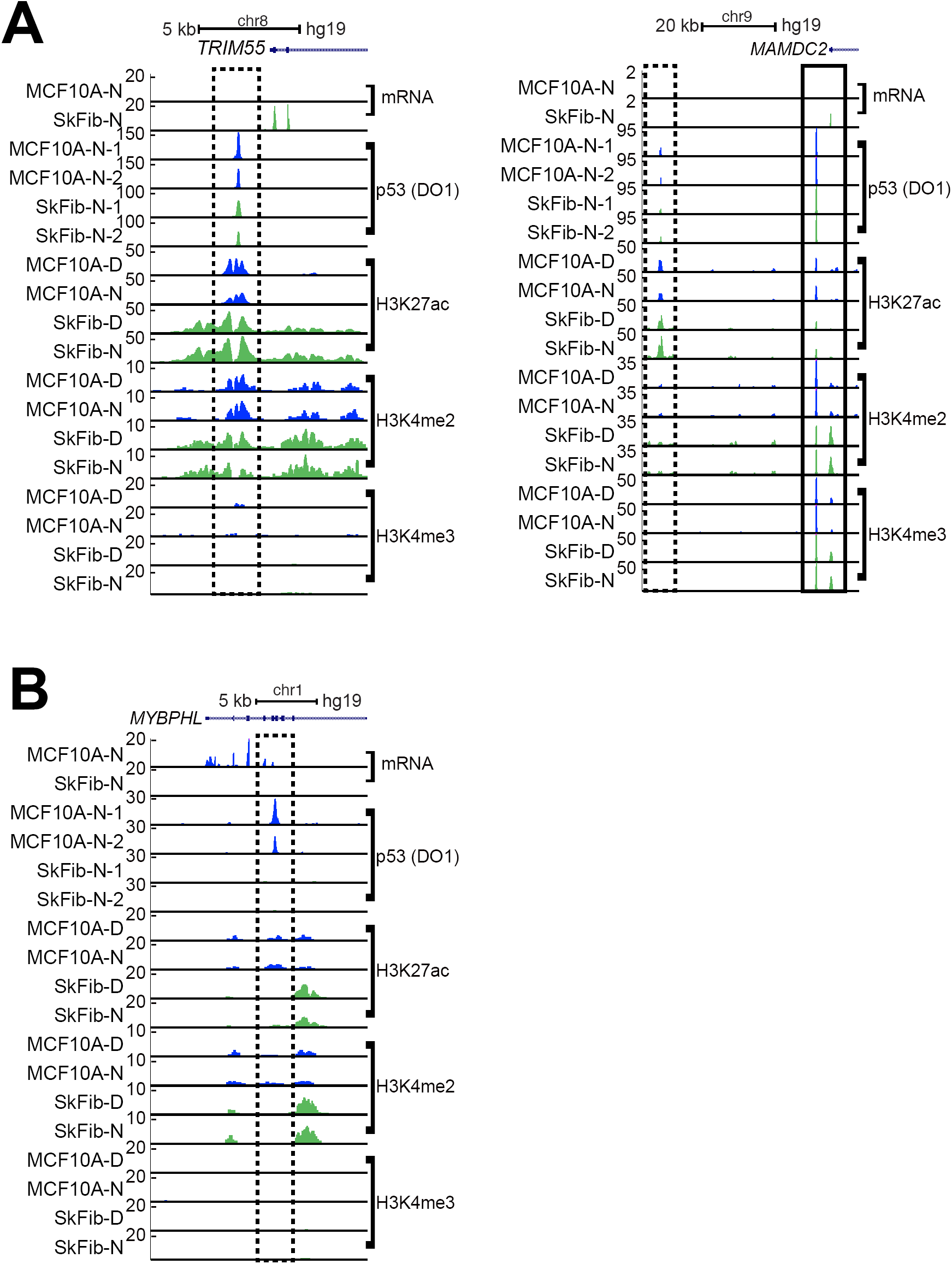
Differential promoter and enhancer activity in cell type-dependent expressed p53 target genes. (A) Representative UCSC Genome Browser track view of TRIM55 and MAMDC2 loci, two fibroblast-specific p53 targets, in response to DMSO (D) or Nutlin-3A (N) treatment. p53- bound, putative enhancers (H3K27ac+, H3K4me2+, H3K4me3-) illustrated by dashed box for TRIM55 and MAMDC2, while the putative MAMDC2 promoter (H3K27ac/H3K4me2/ H3K4me3+) is represented by a solid box. MCF10A-N-1 is biological replicate of MCF10A-N-2, and SkFib-N-1 is biological replicate of SkFib-N-2. (B) Representative UCSC Genome Browser track view at the MYBPHL locus, demonstrating an MCF10A-specific enhancer signature (H3K27ac+,H3K4me2+, H3K4me3-) at a p53 peak in response to DMSO (D) or Nutlin-3A (N) treatment. The y-axis is scaled to the maximum intensity between MCF10A or SkFib for each feature. MCF10A-N-1 is biological replicate of MCF10A-N-2, and SkFib-N-1 is biological replicate of SkFib-N-2.

**Supplemental Figure S3.**
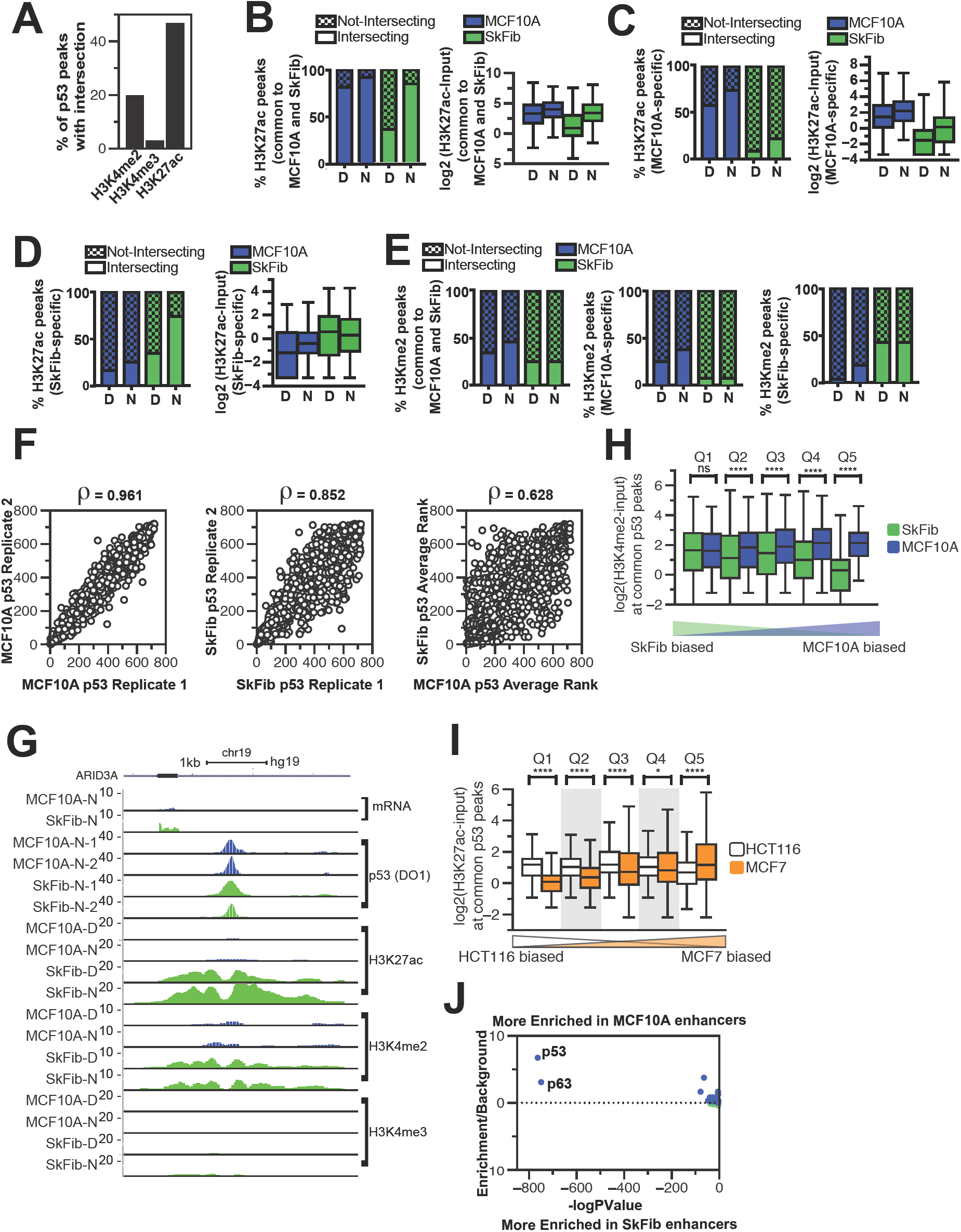
Epigenomic analysis of cell type-dependent p53 activation. (A) The percentage of MCF10A p53 binding sites overlapping H3K4me2, H3K4me3, or H3K27ac under basal (DMSO-treated) conditions. (B-D) The percentage of intersecting H3K27ac peaks (input-normalized, MACS v2, p<0.01) with p53 peaks (input-normalized, MACS v2, p<0.01) that are (B) common to MCF10A and SkFib, (C) MCF10A-specific, or (D) SkFib-specific in response to DMSO (D) or Nutlin-3A (N) treatment. Adjacent boxplots depict H3K27ac enrichment over a 500bp window from the p53-binding site. (E) The percentage of intersecting H3K4me3 peaks (input-normalized, MACS v2, p<0.01) with p53 peaks (input-normalized, MACS v2, p<0.01) that are common to MCF10A and SkFib (left), MCF10A-specific (middle), or SkFib-specific (right) in response to DMSO (D) or Nutlin-3A (N) treatment. (F) Dot plots depicting correlation between rank-ordered biological replicates for MCF10A (left), SkFib (middle), or between MCF10A and SkFib (right). Correlation value (ρ) is Spearman’s rho. (G) Representative UCSC Genome Browser track view at the ARID3A locus, demonstrating similar p53 occupancy between MCF10A and SkFib after Nutlin-3A treatment. The y-axis is scaled to the maximum intensity between MCF10A or SkFib for each feature. MCF10A-N-1 is biological replicate of MCF10A-N-2, and Sk- Fib-N-1 is biological replicate of SkFib-N-2. (I) (F) H3K27ac enrichment in a 500bp window from the peak center (input-subtracted, log2) at common rank-ordered HCT116 or MCF7 p53 binding sites. ****p<0.0001 and *p<0.05, as calculated by a paired Wilcoxon rank-sum test. (J) Homer-derived transcription factor motif enrichment found in MCF10A (top) or SkFib (bottom) specific enhancers (H3K4me2+/H3K27ac+/H3K4me3-). Full list of transcription factor enrichment and facet-specific enhancers are found in Supplemental Table S4.

**Supplemental Figure S4.**
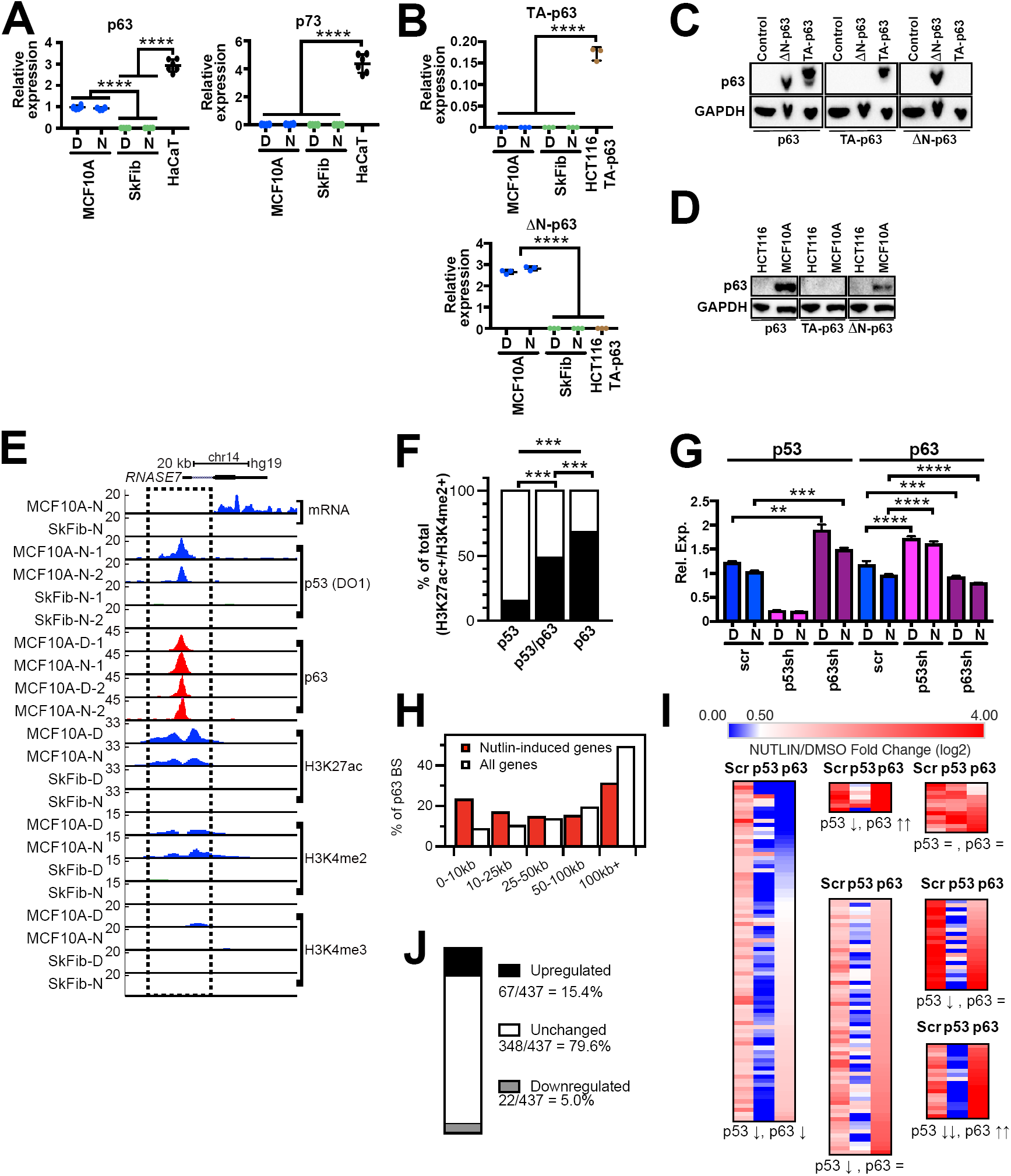
Epithelial-specific p53 binding sites correlate with p63 binding and epithelial-specific enhancers. (A) qRT-PCR analysis of p53, p63 or p73 of cell lysates from MCF10A, SkFib, HUVEC and HaCaT cells at 6 hours after DMSO (D) or Nutlin-3A (N) treatment. Expressions detected by qRT-PCR are normalized to GAPDH expression. HaCaT cells used as positive control for p63 and p73 expression. Error bars represent SEM, p values were calculated by Student’s t-test, ****p<0.0001. (B) qRT-PCR analysis of TA-p63 and ΔN-p63 of cell lysates from MCF10A, SkFib and HCT116-TA-p63 cells. Transiently transfected HCT116 cells with TA-p63 used as a positive control for TA-p63 expression. Error bars represent SEM, p values were calculated by Student’s t-test, ****p<0.0001. (C) Imnunoblotting for p63, TA-p63 and ΔN-p63 of cell lysates from HCT116, HCT116-TA-p63 and HCT116-ΔN transiently transfected cells to confirm antibody specificity for different isoforms. (D) Immunoblotting for p63, TA-p63 and ΔN-p63 of cell lysates from MCF10A cells. HCT116 cells used as negative control. (E) Representative UCSC Genome Browser track view of the RNASE7 locus, illustrating a MCF10A-specific putative enhancer bound by p53 and p63 in response to DMSO (D) or Nutlin-3A (N) treatment (H3K27Ac+, H3K4me2+, H3K4me3-; dashed box). The y-axis is scaled to the maximum intensity for each data set. MCF10A-D-1 is biological replicate of MCF10A-D-2, MCF10A-N-1 is biological replicate of MCF10A-N-2, and SkFib-N-1 is biological replicate of SkFib-N-2. (F) The percentage of intersecting H3K27ac/H3K4me2+ peaks (input-normalized, MACS v2, p<0.01) with p53 only, p63 only, or p53/p63 peaks observed in MCF10A cells (input-normalized, MACS v2, p<0.01) (D) The percentage of p63 binding sites observed in MCF10A cells at varying distances to the nearest TSS of all RefSeq genes (white) or Nutlin-3A induced genes (red). (G) qRT-PCR (right) analysis of p53 and p63 in response to p63 knockdown in MCF10A cells stably expressing shRNA to p63 or a non-targeting control shRNA (scr) after 6 hours of DMSO (D) or Nutlin-3A (N) treatment. Target gene expression is normalized to GAPDH for qRT-PCR analysis. Error bars represent SEM, p values were calculated by Student’s t-test, ****p<0.0001, ***p<0.001 and **p<0.01. (H) The percentage of p63 binding sites observed in MCF10A cells at varying distances to the nearest TSS of all RefSeq genes (white) or Nutlin-3A induced genes (red). (I) K-means clustering of Nutlin-3A/DMSO fold-change values of previously identified MCF10A-specific p53 targets in MCF10A cells expressing either control (scr), p53, or p63-targeted shRNA. (J) The number and percent of p53 target genes showing downregulated, upregulated, or unchanged basal mRNA expression in MCF10A-p63KD cell lines.

**Supplemental Figure S5.**
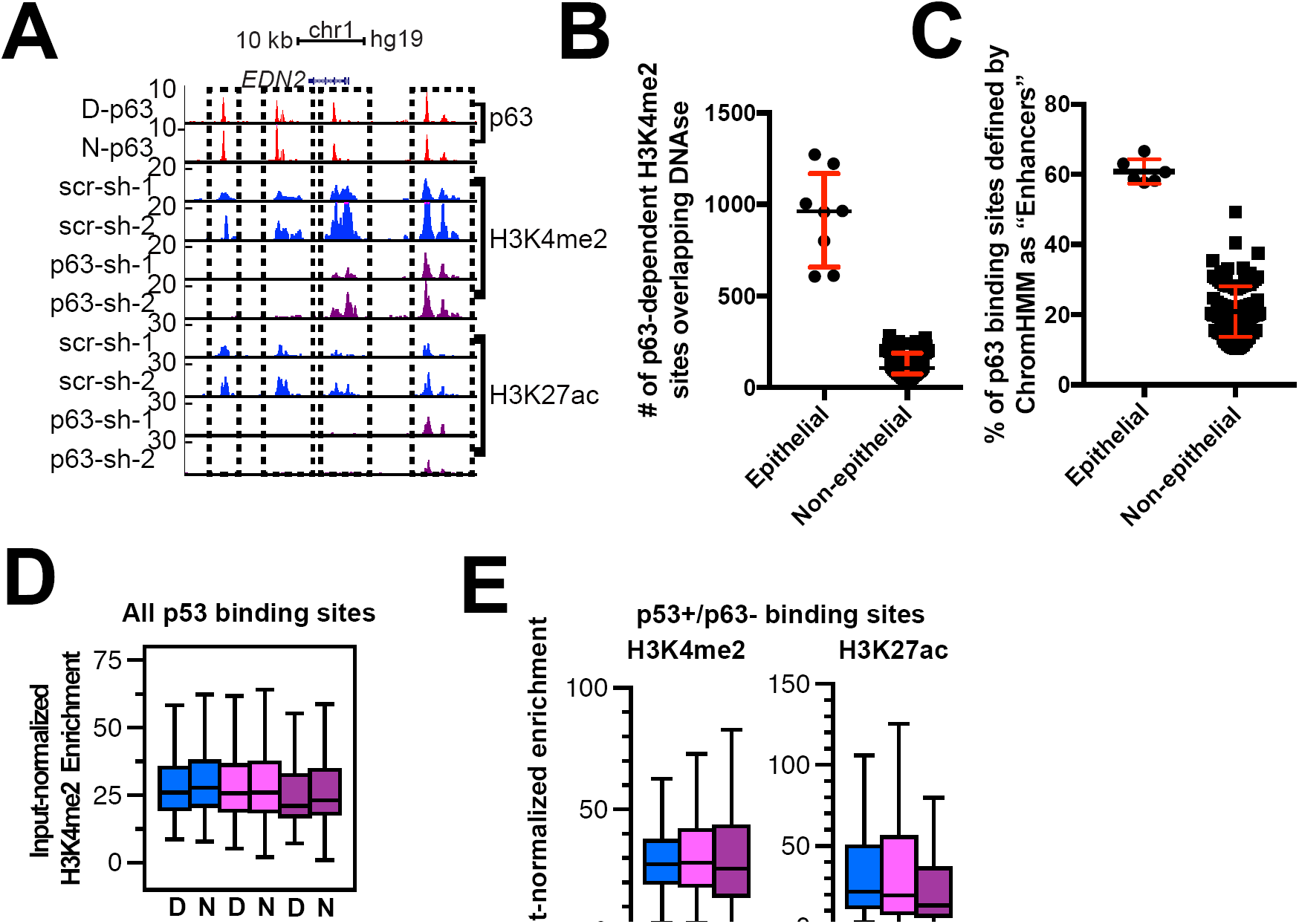
p63 is an enhancer maintenance factor. (A) Representative UCSC Genome Browser track view of the EDN2 locus, illustrating three MCF10A-specific putative enhancers bound by p63 that are lost in response to p63 depletion (H3K- 27ac+, H3K4me2+, H3K4me3-; three separate dashed box). The y-axis is scaled to the maximum intensity for each data set. (B) The number of p63-sensitive, H3K4me2-marked enhancers (out of 1496 total) overlapping DNAse hypersensitive sites (DHS) across epithelial and non-epithelial cell types analyzed by the ENCODE project. Error bars represent the median and 95% confidence interval. (C) The percentage of all MCF10A p63 binding sites intersecting ChromHMM-derived “Enhancer” chromatin states in epithelial and non-epithelial cell types. (D) H3K4me2 enrichment (Input-subtracted H3K4me2, -/+ 250bp from p53 motif center) at p53 binding sites in MCF10A cells expressing control (scr), p53, or p63-targeted shRNA in response to DMSO or Nutlin-3A treatment. (E) Enrichment of H3K4me2 (left) or H3K27ac (right) at MCF10A p53+/p63- binding sites in in MC- F10A cells expressing control (scr), p53, or p63-targeted shRNA in response to DMSO treatment. H3K4me2 and H3K27ac enrichment are reported as input-subtracted histone modification within a -/+ 250bp window from the p53 motif center.

